# Strong regulatory effects of *vgll3* genotype on reproductive axis gene expression in immature male Atlantic salmon

**DOI:** 10.1101/2021.08.08.455548

**Authors:** Ehsan Pashay Ahi, Marion Sinclair-Waters, Jacqueline Moustakas-Verho, Shadi Jansouz, Craig R. Primmer

## Abstract

Age at maturity is a major contributor to the diversity of life history strategies in organisms. The process of maturation is influenced by both genetics and the environment, and includes changes in levels of sex hormones and behavior, but the specific factors leading to variation in maturation timing are not well understood. *gnrh1* regulates the transcription of gonadotropin genes at the onset of puberty in many species, but this gene is lacking in certain teleost species including Atlantic salmon (*Salmo salar*), which raises the possibility of the involvement of other important regulatory factors during this process. Earlier research has reported a strong association of alternative alleles of the *vgll3* gene with maturation timing in Atlantic salmon, suggesting it as a potential candidate regulating reproductive axis genes. Here, we investigated the expression of reproductive axis genes in immature Atlantic salmon males with different *vgll3* genotypes during the spawning period. We detected strong *vgll3* genotype-dependent differential expression of reproductive axis genes (such as *fshb*, *lhb*, *amh* and *igf3*) tested in the pituitary, and testis of one-year-old immature male Atlantic salmon. In addition, we observed differential expression of *jun* (*ap1*) and *nr5a1b* (*sf1*), potential upstream regulators of gonadotropins in the pituitary, as well as *axin2, id3*, *insl3*, *itch*, *ptgs2a* and *ptger4b*, the downstream targets of *amh* and *igf3* in the testis. Hereby, we provide evidence of strong *vgll3* genotype-dependent transcriptional regulation of reproductive axis genes prior to sexual maturation together with models for distinct actions of *vgll3* genotypes on the molecular processes controlling spermatogenesis in Atlantic salmon.

## Introduction

The age at which an individual reproduces, its age at maturity, is a critical time-point in an organism’s life history as it affects fitness traits including survival, and reproductive success (Mobley et al., 2021). There is considerable variation in maturation age both within and among species, and maturation timing thus contributes to the remarkable variation observed in the life history strategies of organisms (Healy et al., 2019). The process of maturation is influenced by both genetics and the environment, and includes changes in levels of sex hormones and behaviour (Varlinskaya et al., 2013), but the specific factors leading to variation in maturation timing are not well understood (Mobley et al., 2021).

The onset of puberty is controlled through the expression of follicle-stimulating hormone (*fshb*) and luteinizing hormone (*lhb*) in the pituitary prior to sexual maturation (Lethimonier et al., 2004). The major upstream regulator of *fshb* and *lhb* is gonadotropin releasing hormone (GnRH), encoded by *gnrh1* gene and expressed in the hypothalamus (Whitlock et al., 2019). Genome of fishes of the family Salmonidae, that includes Atlantic salmon (*Salmo salar*), however, lack *gnrh1* and the compensatory role of other members of the *gnrh* family, *gnrh2* and *gnrh3*, in controlling puberty has remained ambiguous (Whitlock et al., 2019). This suggests the presence of potential upstream regulators of gonadotropin release, other than the *gnrh* family, which can control the transcription of both *fshb* and *lhb* (Jin and Yang, 2014). In Atlantic salmon, the vestigial-like family member 3 gene (*vgll3*) is strongly associated with maturation timing in the wild in both sexes but also exhibits sex-specific maturation effects (Barson et al., 2015; Czorlich et al., 2018). This association between *vgll3* and maturation probability has been subsequently validated in one year-old males in common garden settings (Debes et al., 2021; Sinclair-Waters et al., 2021; Verta et al., 2020). This makes *vgll3* a potential candidate regulating reproductive axis genes in place of the absent *gnrh1*, however, the molecular details of the action of *vgll3* on reproductive axis genes has remained unexplored. It has already been suggested that *vgll3* might affect gametogenesis by regulating cell fate commitment genes (Kjærner-Semb et al., 2018; Kurko et al., 2020), and its repression in salmon testis has been associated with induced onset of puberty (Kjærner-Semb et al., 2018; Verta et al., 2020). A recent study demonstrated that expression of alternative isoforms of *vgll3* associates with variation in age at maturity in one-year-old salmon males (Verta et al., 2020). The *vgll3* alleles previously associated with later and earlier maturation, *vgll3\**L and vgll3*E, respectively, display cis-regulatory expression differences and *vgll3\**L allele expresses a rare transcript isoform in the testis at the pre-pubertal stage. However, the effects of alternative *vgll3* alleles on gene expression of other reproductive axis genes have yet to be investigated.

To further explore the role of *vgll3* in sexual maturation, we examined the expression of reproductive axis genes in immature one year old Atlantic salmon males with different *vgll3* genotypes during the spawning period. We first investigated the expression levels of 9 reproductive axis genes including in the brain, pituitary, and testis of one-year-old immature male Atlantic salmon with different *vgll3* genotypes. Moreover, to elucidate the mechanism by which *vgll3* regulates the expression of reproductive axis genes, we tested the expression level of other potential upstream regulators of *fshb* and *lhb* in the pituitary (Jin and Yang, 2014), and downstream targets of anti-Müllerian hormone (*amh*) and insulin-like growth factor-3 (*igf3*) in the testis (Morais et al., 2017).

## Materials and methods

### Animal material and genotyping

Atlantic salmon used for this study were a subset of individuals used in Sinclair-Waters et al., 2021. Offspring used for this study were selected from a single family resulting from crossing unrelated parents (Kymijoki origin, F1 hatchery generation) with heterozygous (*vgll3*EL*) genotypes, thus enabling assessment of expression patterns of all *vgll3* genotypes within a similar genetic background. Offspring *vgll3* genotypes were determined as described in Sinclair-Waters et al 2021. Sex was determined by internally checking for the presence of female or male gonads during dissection.

### Sample collection, RNA extraction and cDNA synthesis

Tissue sampling took place approximately one-year post-fertilization. Salmon were euthanized with anesthetic overdose (buffered tricaine methanesulfonate (MS-222)) and the brain, pituitary, and testes dissected immediately from sexually immature males (10 *vgll3*EE,* 16 *vgll3*EL*, and 8 *vgll3*LL* individuals) using fine forceps. Tissue samples were snap frozen in liquid nitrogen and stored at −80°C until extraction. Dissected tissues were carefully homogenized using the using a Bullet Blender (Next Advance). We extracted RNA from the sampled brain and testis using the NucleoSpin RNA extraction kit (Biotop), and pituitary using NucleoSpin RNA Clean-up XS (Biotop). The RNA pellets were dissolved in 50 μl of nuclease-free water (Ambion), and genomic DNA removed by incubation with DNaseI. The quality and quantity of RNA were assessed by NanoDrop 1000 v3.7 and Bioanalyzer (Agilent), respectively, and 500 ng of RNA per sample was used for cDNA synthesis using SuperScript™ III Reverse Transcriptase (Invitrogen).

### Candidate genes, designing primers and qPCR

We selected 9 candidate genes with a major role in regulating onset of reproduction, 5 genes upstream of *fshb* and *lhb*, and 6 genes downstream of *amh* and *igf3*, as well as 7 candidate reference genes (Supplementary data). Primer design was conducted as described by Ahi et al., 2019, using the Primer Express 3.0 (Applied Biosystems, CA, USA) and OligoAnalyzer 3.1 (Integrated DNA Technology) (Supplementary data) based on gene sequences retrieved from the annotated *Salmo salar* genome available in the Ensembl database, http://www.ensembl.org. The qPCR reactions were prepared as described by Ahi et al., 2019, using PowerUp SYBR Green Master Mix (Thermo Fischer Scientific), and were performed on a Bio-Rad CFX96 Touch Real Time PCR Detection system (Bio-Rad, Hercules, CA, USA). The details of qPCR program and calculation of primer efficiencies are described by Ahi et al., 2019.

### Analysis of gene expression data

We implemented two common algorithms to validate the most suitable reference gene(s); NormFinder (Andersen et al., 2004) and geNorm (Vandesompele et al., 2002). These algorithms use different analysis approaches to rank the most stably expressed reference genes. The geometric means of the top three ranked most stable reference genes across all individuals were used as normalization factors to calculate the expression levels of our target genes in pituitary (*Hprt1*, *Gapdh* and *Elf1a)* and testis (*Gapdh*, *Actb1* and *Hprt1*) (Table 1). In brain, we used the expression of *Elf1a* as the normalization factor as it was already validated to be the most stable reference gene in this organ in Atlantic salmon (Olsvik et al., 2005). Within each tissue, a biological replicate with the lowest expression level for each gene was selected to calculate ΔΔCq values. The relative expression quantities (RQ values) were calculated by 2^−ΔΔCq^, and their fold changes (logarithmic values of RQs) were used for statistical analysis (Pfaffl, 2001). The student t-test was applied for the direct comparisons of gene expression levels between the genotypes, followed by Benjamini-Hochberg method to correct for multiple comparisons.

**Table 1.** Expression stability ranking of reference genes in testis and pituitary of Atlantic salmon. Abbreviations: NE = Not expressed, SV = stability value, M = mean value of stability.

| Pituitary |  |  |  | Testis |  |  |  |
| --- | --- | --- | --- | --- | --- | --- | --- |
| geNorm |  | NormFinder |  | geNorm |  | NormFinder |  |
| Ranking | M | Ranking | SV | Ranking | M | Ranking | SV |
| *Hprt1* | **0.780** | *Elf1a* | **0.216** | *Gapdh* | **1.021** | *Actb1* | **0.212** |
| *Gapdh* | **0.785** | *Gapdh* | **0.251** | *Actb1* | **1.043** | *Gapdh* | **0.349** |
| *Elf1a* | **0.788** | *Hprt1* | **0.266** | *Hprt1* | **1.107** | *Hprt1* | **0.390** |
| *Rna18s* | 0.834 | *Rna18s* | 0.391 | *Rps20* | 1.117 | *Rps20* | 0.508 |
| *G6pd* | 0.940 | *G6pd* | 0.427 | *Rna18s* | 1.163 | *Rna18s* | 0.522 |
| *Rps20* | 1.050 | *Rps20* | 0.456 | *Ef1a* | 1.183 | *Ef1a* | 0.545 |
| *Actb1* | 1.666 | *Actb1* | 0.625 | *G6pd* | NE | *G6pd* | NE |

## Results

### Expression pattern of reproductive axis genes

There were weak differences between Atlantic salmon individuals with different *vgll3* genotypes in brain expression levels of three *gnrh* family members. *gnrh2* expression was higher in *vgll3*EE* than in *vgll3*EL* and *vgll3*LL* individuals, but only significantly so compared to *vgll3*EL* (Figure 1A). This suggests that the potential regulatory effects of *vgll3* on pituitary expression of gonadotropins may be partly mediated through differential regulation of *gnrh2*, but not *gnrh3a* and *gnrh3b*, which did not show any differences.

**Figure 1.**
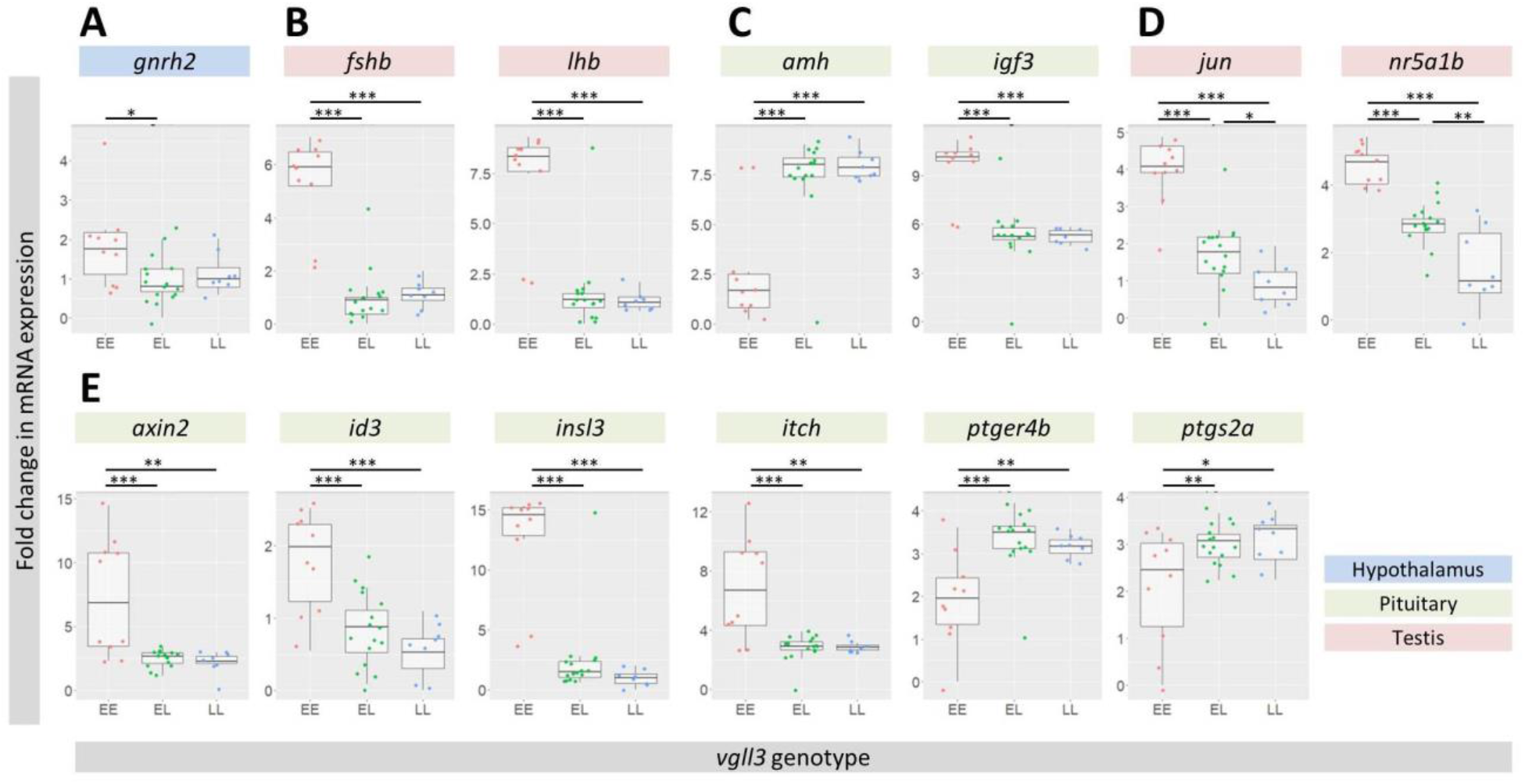
Reproductive axis genes with significant differential expression in juvenile immature male Atlantic salmon with different *vgll3* genotypes. **(A-C)** Major reproductive axis genes with significant differences in expression levels in brain, pituitary and testis. (**D**) Differences in pituitary expression levels of transcription factors at upstream of *fshb* and *lhb*. (**E**) Testis expression levels of downstream targets of *amh* and *igf3*. Small dots indicate individual expression levels, the middle line represents the median, the box indicates the 25/75 percentiles and the whiskers the 5/95 percentiles for each plot. Asterisks above each plot indicate the level of significance for differential expression (* P < 0.05; ** P < 0.01; *** P < 0.001).

The most striking finding of this study was the remarkably higher expression levels of the gonadotropin encoding genes, *fshb* and *lhb*, in the pituitary of *vgll3*EE* individuals compared to the other two *vgll3* genotypes (Figure 1B). This indicates that the induction of gonadotropins has been initiated in the pituitary of *vgll3*EE* males during the reproductive season despite them being immature.

In testis, we observed higher expression of *igf3* and lower expression of *amh* in *vgll3*EE* individuals compared to the two other genotypes (Figure 1C). However, we did not find any differential expression in testis between the genotypes for the receptors of gonadotropins, *fshr* and *lhr* (Supplementary data).

### Expression pattern of selected upstream and downstream genes

We tested the expression of five candidate upstream regulators which have been suggested to bind to promoters of both *fshb* and *lhb* across different groups of vertebrates (Jin and Yang, 2014). Differential expression between *vgll3* genotypes was detected for *jun* (*ap1*) and *nr5a1b* (*sf1*): the highest expression level was seen in *vgll3*EE* individuals, intermediate in *vgll3*EL* and the lowest expression level in *vgll3*LL* (Figure 1D). *vgll3*-related expression differences were not detected for *pitx1* and *foxo1a* (Supplementary data), and the expression levels of *foxp3b* and *nr5a1a* were below the qPCR detection threshold. This suggests a potential direct regulatory link between the *vgll3* alleles and *jun* and *nr5a1b* in the pituitary of male salmon.

We further tested the expression of five known downstream targets of *amh* and a downstream target of *igf3* in testis. The expression level of *vgll3*EE* individuals differed significantly from the other genotypes in all of these genes. *axin2*, a downstream target of *igf3*, showed a similar expression pattern to *igf3* with significantly higher expression in *vgll3*EE* individuals than the other genotypes (Figure 1E). Among the *amh* downstream targets, two genes, *ptgs2a* and *ptger4b*, showed lower, and three genes, *id3*, *insl3* and *itch*, displayed higher expression levels in *vgll3*EE* individuals than the other two genotypes (Figure 1E).

## Discussion

We report the differential expression of reproductive axis genes in the brain (g*nrh2*), pituitary (*fshb*, *lhb*, *jun*, and *nr5a1b*), and testis (*amh*, *igf3*, *axin2*, *id3*, *insl3*, *itch*, *ptger4b*, and *ptgs2a*) in immature one year old Atlantic salmon males with different *vgll3* genotypes. We find significant differences in expression of these genes between *vgll3*EE* individuals compared to *vgll3*EL* and *vgll3*LL* individuals, suggesting that the regulation of these genes is *vgll3* allele-dependent.

The differential expression of *gnrh2* in the brain showed the weakest difference across genotypes, with other *gnrh* genes showing no differences in expression. Although the receptor of *gnrh2* is also expressed in the pituitary of Atlantic salmon during maturation (Ciani et al., 2020), it is uncertain whether the weak changes in *gnrh2* expression can be responsible for the dramatic changes in expression of gonadotropins in this study, or it is likely that other upstream player(s) are involved.

We find higher expression of gonadotropins *fshb* and *lhb* in the pituitary of immature *vgll3*EE* individuals compared to *vgll3*EL* and *vgll3*LL* individuals. Consistent with this, we find that each additional copy of the *vgll3*E* allele increased expression of *jun* and *nr5a1b*, candidate upstream regulators of *fshb* and *lhb* (Jin and Yang, 2014), in the pituitary of immature individuals with *vgll3*EE* individuals having the highest, *vgll3*EL* individuals intermediate, and *vgll3*LL* individuals the lowest relative expression of these genes.

*amh* is a crucial factor in maintaining spermatogonia in an undifferentiated state (Pfennig et al., 2015), whereas upregulation of *igf3* stimulates spermatogonial proliferation and differentiation (Morais et al., 2017). Intriguingly, expression of *amh* was lower and *igf3* higher in the testis of immature *vgll3*EE* individuals as compared to the testis of immature *vgll3*EL* and *vgll3*LL* individuals. This is likely due to higher expression of *fshb* in *vgll3*EE* individuals since *fshb* induces *igf3* and inhibits *amh* expression, which can lead to the onset of maturation (Li et al., 2021). The inhibitory effects of *amh* on spermatogonial differentiation and the stimulatory effects of *igf3* may be mediated by differential regulation of their downstream targets, including *id3*, *insl3*, *itch*, *ptgs2a*, *ptger4b*, and *axin2* in testis.

Insulin-like peptide 3 gene, *inls3*, is expressed in Leydig cells in the testis, and it has been shown in zebrafish that *insl3* promotes spermatogonial differentiation and mediates the stimulatory effect of *fshb* on spermatogenesis (Assis et al., 2016). Moreover, Sertoli cell-derived amh inhibits spermatogonial differentiation partly through reducing the expression levels of *insl3* in Leydig cells in adult zebrafish (Skaar et al., 2011). Consistently, we found reduced expression of *insl3* in testis accompanied with increased expression level of *amh* in *vgll3*LL* individuals of immature Atlantic salmon. This suggests *amh* dependent *insl3* inhibition as a potential molecular mechanism by which spermatogonial differentiation is delayed in *vgll3*LL* individuals.

Another likely scenario for inhibition of spermatogonial differentiation in *vgll3*LL* individuals could be the increased expression of *ptgs2a* and *ptger4b*, which are known to be transcriptionally induced by *amh* in fish testis (Morais et al., 2017). *ptgs2a* encodes a key enzyme involved in production of a prostaglandin (PGE_2_), which reduces the mitotic activity of differentiating spermatogonia, and thereby inhibits spermatogenesis (Crespo et al., 2020). The inhibition of PGE_2_ on the development of spermatogonia seems to be mediated through a receptor encoded by *ptger4b* expressed mainly by testicular somatic cells (Crespo et al., 2020). Taken together, these findings imply that in immature salmon with the *late vgll3* allele, sexual maturation is delayed through an *amh* dependent mechanism and probably at the differentiation stage of spermatogonia (summarized in Figure 2A).

**Figure 2.**
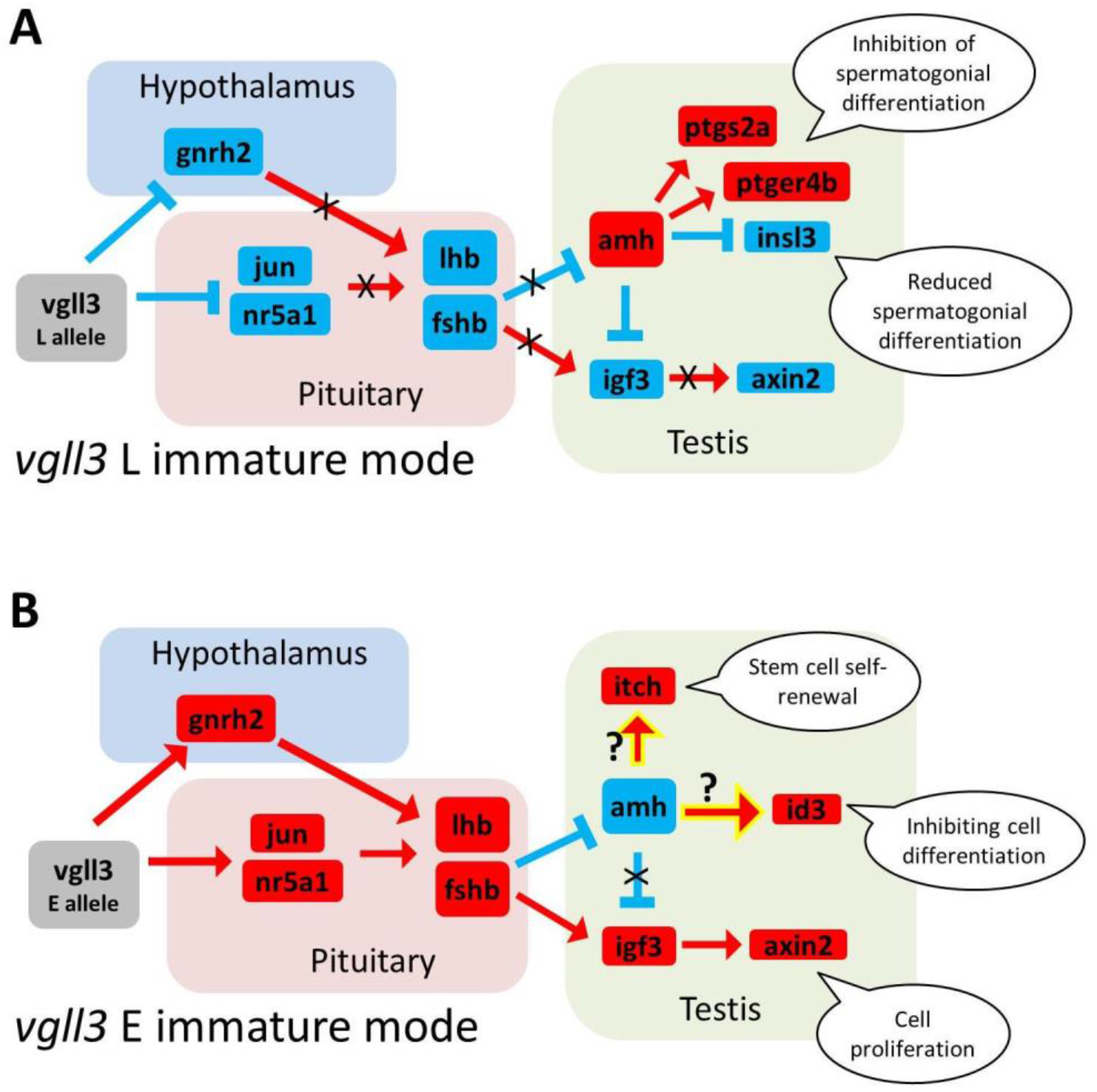
Models for alternative modes of potential *vgll3*-allele dependent regulatory interactions between the differentially expressed genes along the reproductive axis in premature male Atlantic salmon. Red arrows indicate transcriptional induction, blue blocked lines represent transcriptional inhibition and black crosses display expected inhibition of transcriptional regulatory connections, whereas question marks on the highlighted arrows indicate unexpected regulatory outcomes via currently unknown molecular mechanism(s). We hypothesize that the *vgll3*\*L allele promotes the premature state in the testis via inhibition during the final stages of spermatogonia differentiation, whereas the premature state in *vgll3*EE* individuals results from cell fate commitment prevention and thus the maintenance of pluripotency in the testis via an unknown *amh* independent mechanism.

The gene expression results were more complicated to interpret in the immature *vgll3*EE* salmon, since both gonadotropins had much higher expression levels in pituitary than the other genotypes as well as higher expression of *igf3* and lower expression of *amh* in testis. These expression patterns are reminiscent of the mature state (Morais et al., 2017); however, all these individuals, sampled during the spawning period, were immature. This indicates that although these molecular changes were consistent with promotion of maturation, it appears they were not sufficient for inducing maturation in these individuals. In *vgll3*EE* fish, the increased expression of gonadotropins was accompanied by increased expression of *igf3*, a growth factor with gonad specific expression in teleost fish, which is induced by *fshb* and mediates its effects on testis (Li et al., 2021). Moreover, the increased expression of *axin2*, a downstream target of Igf3 involved in proliferation of undifferentiated spermatogonia (Safian et al., 2018), indicates functional activity of Igf3 in the testis of *vgll3*EE* fish. In contrast, the reduced expression of *amh* was not accompanied by similar changes in two of its downstream effectors, *itch* and *id3*, in *vgll3*EE* fish testis. This might explain why these individuals had not entered the maturation stage in the testis, since both *itch* and *id3* are known to inhibit cell fate commitment by controlling the maintenance of self-renewal, pluripotency and the undifferentiated state of progenitor cells (Morais et al., 2017) (summarized in Figure 2B). Why the reduced expression of *amh* did not result in decreased expression of *itch* and *id3* in *vgll3*EE* fish thus requires further investigations. To summarize, the *late vgll3* allele appears to promote the premature state in the testis via inhibition during the final stages of spermatogonia differentiation, whereas the premature state in *vgll3*EE* individuals might be a result of cell fate commitment prevention and thus the maintenance of pluripotency in the testis via an unknown *amh* independent mechanism (Figure 2).

## Acknowledgements

We acknowledge Jaakko Erkinaro and staff at the Natural Resources Institute Finland (Luke) hatchery in Laukaa and members of the Evolution, Conservation and Genomics research group for their help in coordinating and collecting gametes for crosses. We thank Jukka-Pekka Verta and Iikki Donner for valuable discussions. We also thank Nikolai Piavchenko for help with fish husbandry as well as Iikki Donner, Seija Tillanen and Annukka Ruokolainen for laboratory assistance.

## Author Contributions

EP, CRP, MS-W, JM, SJ conceived the study; EP, MS-W, JM, SJ performed experiments; EP developed methodology and analyzed the data; EP, CRP, MS-W, JM interpreted results of the experiments; EP, CRP, MS-W, JM drafted the manuscript, with EP having the main contribution; SJ edited manuscript and all authors approved the final version of manuscript.

## Ethical approval

Animal experimentation followed European Union Directive 2010/63/EU under license ESAVI/42575/2019 granted by the Animal Experiment Board in Finland (ELLA).

## Data availability

Supplementary file containing raw expression data and primers information

